# Comparison of sporulation and germination conditions for *Clostridium perfringens* strains

**DOI:** 10.1101/2023.02.16.528852

**Authors:** Marc Liggins, Norma Ramírez Ramírez, Ernesto Abel-Santos

**Author notes:** **Contact:** 4505 Maryland Parkway, Campus Box 4003, Las Vegas, NV 89154.

## Abstract

*Clostridium perfringens* is a spore forming, obligate anaerobe, Gram-positive bacterium that causes a range of diseases in humans and animals. The infectious form of *C. perfringens* is the spore, a structure that is derived from the vegetative cell under conditions of nutrient deprivation. The first step in *C. perfringens* pathogenesis is the differentiation of spores into replicating bacteria. Previous work in analyzing *C. perfringens* spore germination has produced contradictory, strain-specific results. Hence, we analyzed the requirements for spore formation and germination in seven different *C. perfringens* strains. Our data showed that *C. perfringens* sporulation conditions are strain specific, but germination responses are homogenous. *C. perfringens* spores can germinate using two distinct pathways. The first germination pathway (the amino acid-only pathway or AA) requires L-alanine, L-phenylalanine, and sodium ions (Na^+^) as co-germinants. L-arginine is not a required germinant but potentiates germination. The AA pathway is inhibited by aromatic amino acids and potassium ions (K^+^). Bicarbonate (HCO3-), on the other hand, bypasses potassium-mediated inhibition of *C. perfringens* spore germination through the AA pathway. The second germination pathway (the bile salt and amino acid pathway or BA) is more promiscuous and is activated by several bile salts and amino acids. In contrast to the AA pathway, the BA pathway is insensitive to Na^+^, but can be activated by either K^+^ or HCO3-. We hypothesize that *C. perfringens* may have evolved these two distinct germination pathways to ensure spore response to different host environments.

**Manuscript contribution to the field:** *Clostridium perfringens* is a ubiquitous bacterium that can infect a wide variety of animal hosts, including humans. *C. perfringens* counts with a veritable arsenal of toxins that are differentially expressed depending on the host infected. Besides toxin production, *C. perfringens* can also form dormant and resistant spores that serve as infective vehicles. *C. perfringens* spores need to germinate back into vegetative cells to cause disease. Befitting to its wide host range, *C. perfringens* spore germination seems to use strain specific. In this study, we tested the ability of seven *C. perfringens* strains to produce spores under different conditions. We also tested the requirements for spore germination. We found that while *C. perfringens* sporulation was highly varied, the germination response for all strains could be divided into two distinct pathways. Since *C. perfringens* spores need to germinate to cause infection, understanding the germination behavior could lead to approaches for the prevention of diseases in humans and veterinary animals.

## Background

*Clostridium perfringens* is a gram-positive, rod-shaped, spore-forming, obligate anaerobic bacterium [1, 2]. *C. perfringens* is a very versatile pathogen that has adapted to a number of different environments and hosts. This ability allows *C. perfringens* to causes a wide range of diseases in humans and animals [3].

The ability of *C. perfringens* to cause disease in different hosts is ascribed mainly to the differential production of four major and ten minor protein toxins. The most important toxins linked to infections are the alpha toxin (CPA), beta toxin (CPB), enterotoxin (CPE), epsilon toxin (ETX), iota toxin (ITX), and necrotic enteritis toxin B-like (NetB) [4].

*C. perfringens* are classified into 7 toxinotypes based on their ability to produce different toxins and infect different hosts [5]. *C. perfringens* type A produces CPA as the main determinant for myonecrosis (gas gangrene), while also producing CPE and NetB during gastrointestinal infections. *C. perfringens* type B targets mostly sheep while producing CPA, CPB, and ETX. *C. perfringens* type C causes necrotic enteritis in neonatal hosts and produces both CPA and CPB. *C. perfringens* type D causes enterotoxemia in Caprinae species using CPA and ETX. Finally, *C. perfringens* type E is not strongly associated with mammalian infections but can produce both CPE and ITX [6, 7]

When stressed, *C. perfringens* forms resistant and dormant spores [8]. *C. perfringens* spores persist in the environment and serve as infective vehicles [9]. To start an infection, *C. perfringens* spores need to germinate into toxin-producing vegetative cells [10].

Commonly, the first step to trigger spore germination is the detection of nutrients by a family of membrane proteins (Ger receptors) [10, 11]. Most Ger receptors are encoded in tricistronic operons that produce three distinct protein subunits [12]. Most *Bacilli* and *Clostridia* species encode multiple Ger receptors that allows for the recognition of different germination signals. In contrast, *C. perfringens* encodes a single functional receptor (GerK) even though the spores must be able to germinate in very different environments to cause each tissue-specific infection [13].

As befitted for such a versatile pathogen, *C. perfringens* spores have been reported to recognize different sets of germinants. Parades-Sabja *et al*. reported that all *C. perfringens* spores tested use KCl and L-asparagine as a universal germination mixture [14]. Additionally, *C. perfringens* strains carrying a chromosomal enterotoxin (*cpe*) gene can also germinate with KCl alone. In contrast, *C. perfringens* strains carrying a plasmid-borne *cpe* gene can germinate with L-alanine or L-valine, in addition to KCl/L-asparagine [14]. These results are intriguing. Firstly, chromosomal enterotoxin (from SM101 strain) and plasmid enterotoxin (from F5603 strain) are 100% identical at the amino acid level. Secondly, Although *cpe* is expressed during sporulation, the enterotoxin remains in the mother cell and not in the spore [15]. Thus, the function of the enterotoxin in modulating germination response of *C. perfringens* spores is not clear.

Other studies have expanded the potential germinant diversity used by *C. perfringens* spores. Wax and Frees reported that *C. perfringens* spores were also able to germinate using L-asparagine, D-glucose, D-fructose, and potassium ions (AGFK), a mixture that is commonly used to induce *B. subtilis* spore germination [16]. A separate study found that sodium ions and phosphate instead of KCl acted as germinants of *C. perfringens* spores, but only in food poisoning isolates [17]. In contrast, Kato *et al*. showed that bicarbonate, L-alanine, and inosine were capable of inducing *C. perfringens* spore germination, a response that closely mimics the germination profile of *B. cereus* and *B. anthracis* spores [18].

The unorthodox germination behavior of *C. perfringens* spores is further highlighted by the fact that all strains sequenced encode a single, identical GerK receptor [13]. Thus, if strain-specific germination responses are dependent on GerK activation, this receptor must be able to recognize structurally different molecules. Furthermore, each *C. perfringens* strain uses only a subset of available co-germinants. Hence, the GerK receptor must have multiple binding sites to be able to change substrate specificity between strains. Alternatively, some of the germinants could be recognized by unidentified germination receptors. [19].

Differences in *C. perfringens* spore germination could be potentially attributed to dissimilar assay conditions. Indeed, previous works in *Bacillus* species have shown that sporulation environments can subsequently affect spore germination [20-22]. Another possibility is that spores from different *C. perfringens* strain could recognize unique sets of germinants.

Indeed, inter-strain variability in germination response has been shown in *B. cereus* spores [23, 24]. The heterogeneity of spore germination can be important for *C. perfringens* since each pathogenic strain may infect a limited number of hosts [25].

In this study, we systematically tested sporulation conditions, strain-specificity, and germinant mixtures as variables of *C. perfringens* spore germination. Our results show that each *C. perfringens* strain sporulates under unique conditions, but spore germination was conserved among all strains tested. Contrary to previous results, neither asparagine nor phosphate nor inosine were necessary for spore germination. Instead, spores from all *C. perfringens* strains tested germinated using two distinct pathways. In the first pathway (the amino acid-only or AA germination pathway), *C. perfringens* spores need L-phenylalanine and L-alanine as germinants. The second germination pathway (the bile salt/amino acid or BA germination pathway) is activated by mixtures of bile salts and amino acids. We also showed that sodium and bicarbonate differentially regulate *C. perfringens* spore germination through both pathways. Finally, we found that aromatic amino acids and potassium inhibited the AA germination pathway but acted as a germinants in the BA germination pathway.

## Methods

### Materials

All chemicals were purchased from Sigma-Aldrich Corporation (St. Louis, MO). Thioglycollate medium, peptones, yeast extract, raffinose and agar were purchased from VWR (West Chester, PA). *C. perfringens* strains JGS1936, JGS1473, JGS1882, JGS1521, JGS4104, JGS4151, and JGS 4064 [26] were a generous gift from Prof. J. Glenn Songer (Iowa State University, Ames, IA). The identities of selected *C. perfringens* spore preparations were confirmed by 16S RNA sequencing.

### Testing of growth conditions on *C. perfringens* sporulation yields

*C. perfringens* strains were plated on 2% agar supplemented with 1% yeast extract, 0.1% sodium thioglycollate, 1.5% protease peptone, and 60 mM Na2HPO4. Plates were incubated overnight in an anaerobic environment (5% CO2, 5% H2, 90% N_2_). Single-cell clones were picked and grown for four hours in either thioglycollate medium or BHI broth. All *C. perfringens* strains were then plated on 2% agar supplemented with 1% yeast extract, 0.1% sodium thioglycollate, 60 mM Na_2_HPO_4_ and 1.5% of a peptone source (protease peptone #1, protease peptone #2, protease peptone #3, or potato peptone). Media were also supplemented with 0.5% of a filter-sterilized carbon source (glucose, starch, or raffinose). Some plates were supplemented with theobromine to 0.01% final concentration. Plates were incubated for up to14 days at 37 °C under anaerobic conditions.

Sporulation was quantified by microscopy observation of culture samples stained using the Schaeffer-Fulton method [27]. Under these conditions, spores are stained green and vegetative cells are stained red. The approximate number of green spores and red vegetative cells were counted in at least three independent microscopy fields selected at random. High level of sporulation was defined as >40% spores. Medium level of sporulation was defined as 20-40% spores. Low level of sporulation was defined as <20% spores.

### Purification of *C. perfringens* spores

Each *C. perfringens* strain was plated under their best sporulation conditions (Table S2). Plates were incubated for 5-10 days at 37 °C in an anaerobic environment. The resulting bacterial lawns were collected by flooding with ice-cold deionized water. Spores were pelleted by centrifugation and resuspended in fresh deionized water. After two washing steps, spores were separated from vegetative and partially sporulated cells by centrifugation through a 20%-50% HistoDenz gradient. Spore pellets were washed five times with water, resuspended in 0.1% sodium thioglycollate and stored at 4 °C. All spore preparations were more than 95% pure as determined by microscopy observation of Schaeffer-Fulton stained aliquots.

### Preparation of germinant solution

AGFK mixture (10 mM L-asparagine, 10 mM D-glucose, 10 mM D-fructose, 50 mM KCl) was prepared as previously described [16]. The defined medium employed was described previously [28]. Briefly, a buffer solution was made with 6.6 mM KH2PO4, 15 mM NaCl, 59.5 mM NaHCO_3_ and 35.2 mM Na_2_HPO_4_. Three solutions were prepared in using this buffer as diluent. The first solution contained all salts at 1000X concentrations (final concentration were 10 mg/l MgSO_4_•7H_2_O, 5 mg/l FeSO_4_ •7H_2_O, 5 mg/l MnCl_2_•4H_2_O). The second solution contained vitamins at 10X concentrations (final concentrations were 0.05 mg/l D-biotin, 0.1 mg/l *p*-amino benzoic acid, 0.05 mg/l thiamine hydrochloride, 0.05 mg/l pyridoxine, 1.0 mg/l nicotinic acid). The third solution contained all amino acids, except cysteine at 10X (final concentrations were 10 mM for each amino acid). Cysteine was prepared separately as a 10X solution in 0.2 N HCl. To prepare the defined medium, different solutions were added to buffer at the final concentrations indicated. In some samples, inosine was added to 1 mM final concentration.

To determine individual germinants, stock (10X) solutions of L-amino acids, NaHCO3, KHCO3, KCl, KBr, NaCl, NaBr, and bile salts were individually prepared in deionized sterile water. Combinations of these solutions were tested to determine germinants necessary for *C. perfringens* spore germination.

### Requirements for *C. perfringens* spore germination

Changes in light diffraction during spore germination were monitored at 580 nm (OD_580_) on a Tecan Infinite M200 96-well plate reader (Tecan group, Männedorf, Switzerland). *C. perfringens* spores were heat-activated at 65 °C for 30 min [29]. The spore suspension was cooled to room temperature and monitored for auto-germination for 30 min. Germination experiments were carried out with spores that did not auto-germinate. After heat activation, spores were resuspended to an OD_580_ of 1 in AGFK, LB broth, or defined medium. Spore germination rates were evaluated based on the decrease in OD_580_ at room temperature. After germinant additions, OD_580_ was measured at 1 minute intervals for 90 minutes. Relative OD_580_ values were derived by dividing each OD_580_ reading by the initial OD_580_. Experiments were performed in triplicate with at least two different spore preparations. Germination rates were calculated from the initial linear region of the germination curves. Standard deviations were calculated from at least six independent measurements and were typically below 20%. Germination was confirmed in selected samples by microscopy observation of Schaeffer-Fulton stained aliquots.

To determine amino acid co-germinants, *C. perfringens* spores were resuspended in germination buffer (0.1 mM sodium phosphate buffer (pH 6.5), 50 mM NaHCO_3_) to an OD_580_ of 1. Putative germinants were added individually or in combinations to a final concentration of 10 mM. After addition of germinants, spore germination was monitored by the decrease in optical density at 580 nm, as above. Germination rates were set to 100% for *C. perfringens* spores germinated in the presence of L-alanine and L-phenylalanine. Relative germination for other germinant combinations was calculated as the fraction of germination rate compared to germination with L-alanine/L-phenylalanine.

To determine bile salt co-germinants, *C. perfringens* spores were resuspended in potassium phosphate buffer (pH 6.5) supplemented with 5% KHCO3, and 150 mM KCl. Spore germination was started by addition of 6 mM taurocholate, and 6 mM individual amino acids. *C. perfringens* spores were also germinated with 6 mM L-alanine and 6 mM individual bile salts.

After addition of germinants, spore germination was monitored as above. Germination rates were set to 100% for *C. perfringens* spores germinated in the presence of L-alanine and taurocholate. Relative germination for other germinant combinations was calculated as the fraction of germination rate compared to germination with L-alanine/taurocholate.

### Testing for inhibitors of *C. perfringens* spore germination

*C. perfringens* spores were resuspended in sodium phosphate buffer (pH 6.5) supplemented with 5% NaHCO3, and 150 mM NaCl (for the AA pathway) or potassium phosphate buffer (pH 6.5) supplemented with 5% KHCO3, and 150 mM KCl (for the BA pathway). Spores samples were then individually supplemented with 6 mM amino acid or 6 mM bile salt analogs. Spore suspensions were incubated for 15 min at room temperature while the OD_580_ was monitored. If no germination was detected, spores were supplemented with 6 mM L-alanine/6 mM L-phenylalanine (for the AA pathway) or 6 mM L-alanine/6 mM taurocholate (for the BA pathway). Germination rates were set to 100% for *C. perfringens* spores germinated in the absence of inhibitor. Relative germination for conditions was calculated as the fraction of germination rate compared to no inhibitor.

### Effect of buffer and pH on *C. perfringens* spore germination

Individual *C. perfringens* spore aliquots were individually resuspended in 0.1 M sodium phosphate buffer (or 0.1 M potassium phosphate buffer and pH values were individually adjusted between 5.5 and 8.0. Germination was started by addition of 6 mM L-alanine/6 mM L-phenylalanine (for the AA pathway) or 6 mM L-alanine/6 mM taurocholate (for the BA pathway). Spore germination was monitored as above. For the AA pathway, the germination rate was set to 100% for *C. perfringens* spores germinated at pH 6.5 in sodium phosphate buffer. For the BA pathway, the germination rate was set to 100% for *C. perfringens* spores germinated at pH 6.5 in potassium phosphate buffer. The percentage of germination for other conditions was calculated as a fraction of the rate of germination at pH 6.5.

### Effect of cations and anions on *C. perfringens* spore germination

*C. perfringens* spores were individually incubated for five minutes in 0.1 M sodium phosphate buffer, pH 6.5 or 0.1 M potassium phosphate buffer, pH 6.5. Samples were then individually supplemented with 150 mM KCl, KBr, NaCl, NaBr, KHCO_3_ or NaHCO_3_. Germination was started by addition of 6 mM L-alanine/6 mM L-phenylalanine (for the AA pathway) or 6 mM L-alanine/6 mM taurocholate (for the BA pathway). Spore germination was monitored by the decrease in optical density, as above. For the AA pathway, the germination rate was set to 100% for *C. perfringens* spores germinated in sodium phosphate buffer without added salts. For the BA pathway, the germination rate was set to 100% for *C. perfringens* spores germinated in potassium phosphate buffer without added salts. The percentage of germination for other conditions was calculated as a fraction of the rate in the absence of added salts.

## Results

*Clostridium perfringens* have shown a lot of variability in both sporulation [30-32] and germination responses [14, 17, 18]. To determine whether these sporulation and germination differences correlated to host specificity, seven different *C. perfringens* strains were tested (Tabel A1). These strains were obtained from healthy and diseased avian and mammalian animals [26, 33].

### Sporulation of *C. perfringens* strains

Previous works have shown that sporulation conditions can affect the subsequent germination response of bacterial spores [20, 22]. To test if *C. perfringens* spore germination could be modulated by sporulation media, we created a matrix of conditions for sporulation with combinations of different liquid media, solid media, carbon sources, peptones, and additives for every *C. perfringens* strain used in this study (Table S1). All *C. perfringens* strains tested sporulated in solid media, but not in liquid media.

However, it was observed that sporulation was dependent upon which liquid media was used for overnight growth prior to plating in agar. For strains JG1936, JG1882, JG4064, overnight growth in BHI was necessary to induce sporulation upon replating in their preferred solid media. Other strains (JGS1521, JGS4151) required overnight growth in liquid thioglycollate medium to induce sporulation in agar. For other strains (JGS1473, JGS4104), the liquid media used for overnight growth changed the preference of solid media required for sporulation.

Glucose, starch, and raffinose were tested as carbon sources for sporulation. Consistent with prior results, raffinose was the preferred carbon source for *C. perfringens* sporulation [30]. In our hands, glucose and starch induced poor sporulation under all conditions tested [32].

Peptone sources have been previously shown to affect the level of sporulation in *C. perfringens* strains [31]. Peptone protease #1 induced sporulation in strains JGS1936, JGS1473, JGS4064, and JGS1521. Peptone protease #2 was able to induce sporulation in strains JGS1882 and JGS4151. Peptone protease #3 induced sporulation in JGS1473 and JGS4104. Potato peptone can induce high levels of sporulation in some *C. perfringens* strains [31]. In our hands, however, potato peptone did not induce sporulation in any of the strains tested.

Theobromine has been reported to increase the levels of sporulation in *C. perfringens* strains [30]. Indeed, strains JGS4104, JGS4064, JGS1521, and JGS4151 only sporulated robustly when theobromine was added to solid media. Strains JGS1936, JGS1882, and JGS1473 sporulated in the absence of theobromine. In the presence of theobromine, sporulation levels for strain JGS1936 remained unchanged, increased for strain JGS1882, and decreased for strain JGS1473.

Aside from differentially affecting sporulation levels, theobromine also reduced sporulation times in strains JGS1936, JGS1473, and JGS1882. In the absence of theobromine, sporulation was not detected until five days post-plating and maximum sporulation level was achieved 7-14 days post-plating. In the presence of theobromine, spores could be detected two days after plating and maximum sporulation levels were seen 5-7 days post-plating.

### Identification of amino acids and sterane germinants of *C. perfringens* spores

Contrary to previous work, *C. pefringens* spores failed to germinate with AGFK, KCL/L-asparagine, sodium/phosphate, or L-alanine/inosine mixtures [14, 17, 18]. Like other *Clostridium* species, *C. perfringens* spores germinated efficiently in defined medium (Fig. 1A). *C. perfringens* spores germinated at the same rate in defined medium containing only amino acids. Henceforth, we refer to this germination response as the amino acid-only (AA) germination pathway.

**Figure 1.**
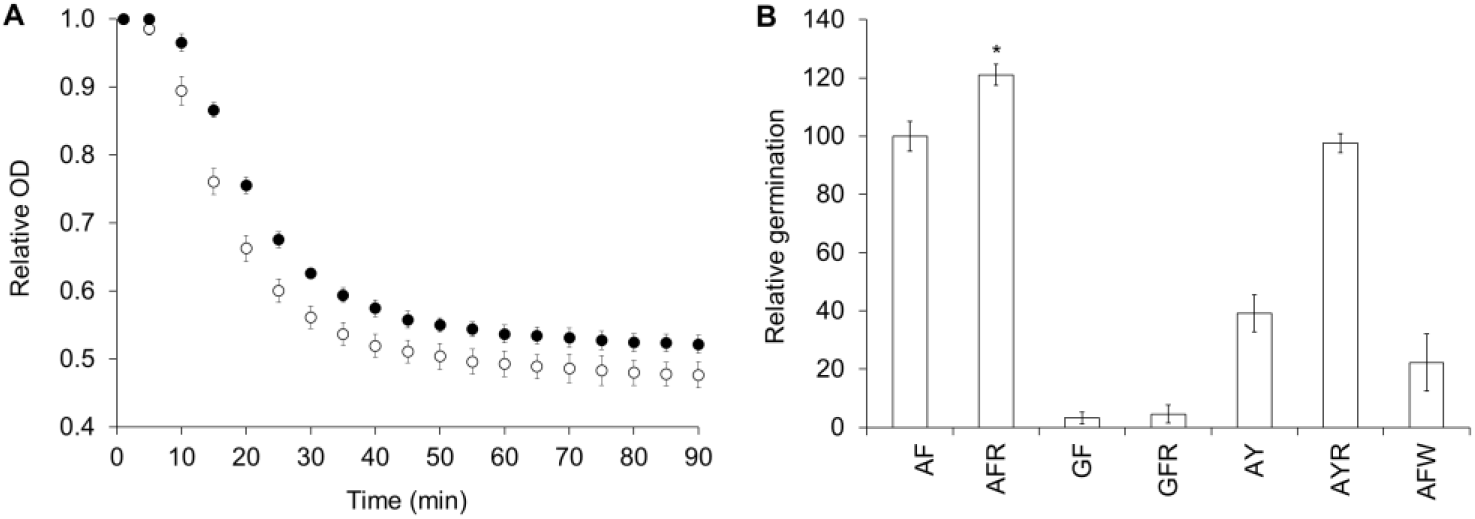
L-alanine/L-phenylalanine-mediated C. perfringens spores germination is potentiated by L-arginine and inhibited by L-tryptophan. (A). *C. perfringens* spores were germinated in defined medium (○) or in a solution containing 25 mM L-alanine, 5 mM L-phenylalanine and 50 mM NaHCO3 (●). Germination was followed by the decrease of optical density at 580 nm (OD580). For clarity, data are shown at 5-min intervals. Data corresponds to *C. perfringens* strain JGS 1936 spores Other *C. perfringens* strains yielded similar results. (B) *C. perfringens* JGS 1936 spores were treated with amino acid mixtures. Germination rates were calculated from the linear segment of optical density changes over time. Relative germination was calculated as fractions of germination rate for spores treated with L-alanine/L-phenylalanine. Amino acids are represented by one-letter code. Error bars represents standard deviations of six independent measurements. * p<0.003 compared to L-alanine/L-phenylalanine.

To identify which amino acids are required for germination, *C. perfringens* spores were exposed to mixtures of small (L-Ala and Gly), polar (L-Ser, L-Thr, and L-Cys), hydrophobic (L-Leu, L-Ile, L-Met, and L-Val), aromatic (L-Phe, L-Tyr, and L-Trp), basic (L-Arg, L-Lys, and L-His), acidic (L-Asp and L-Glu), amide (L-Asn and L-Gln), or constrained (L-Pro) amino acids. None of these solutions alone was sufficient to trigger spore germination. *C. perfringens* spores were then resuspended in solutions containing pairs and trios of the above amino acid groups. *C. perfringens* spore germination was only observed in solutions containing mixtures of small and aromatic amino acids. Faster *C. perfringens* spore germination rates were observed when small and aromatic amino acids were supplemented with basic amino acids.

To further narrow the identity of L-amino acid germinants, all possible combinations of small, aromatic, and basic amino acids were tested individually for their effect on *C. perfringens* spore germination. For all strains tested, strong *C. perfringens* spore germination was seen in the presence of L-alanine/L-phenylalanine (Fig. 1A). L-arginine was not required to trigger germination, but increased germination rates by 20% (Fig. 1B). In the L-alanine/L-phenylalanine germination mixture, L-alanine could not be substituted for glycine, even in the presence of L-arginine. L-tyrosine can substitute L-phenylalanine, but the germination rate was more than 50% slower. Addition of L-arginine to L-alanine/L-tyrosine-treated spores increased germination rates more than 2-fold. *C. perfringens* spores did not respond to L-alanine/L-tryptophan mixtures. In fact, L-tryptophan behaved as an inhibitor of L-alanine/L-phenylalanine-mediated *C. perfringens* spore germination (Fig. 1B).

We recently showed that sterane compounds can modulate the germination response of *C. difficile* and *C. sordellii* spores [34]. Taurocholate, a known co-germinant of *C. difficile* spores, was not sufficient to induce germination in *C. perfringens* spores. On the other hand, combinations of taurocholate and a variety of amino acids induced strong *C. perfringens* spore germination. In fact, only six amino acids did not synergize with taurocholate to induce significant *C. perfringens* spore germination (Fig. 2A). Glycocholate, taurochenodeoxycholate, and taurodeoxycholate also induced *C. perfringens* spore germination in the presence of amino acids. Cholate, chenodeoxycholate, and deoxycholate did not induce nor inhibited *C. perfringens* spore germination in the presence of L-alanine (Fig. 2B). Henceforth, we refer to this germination response as the bile salt/amino acid (BA) germination pathway.

**Figure 2.**
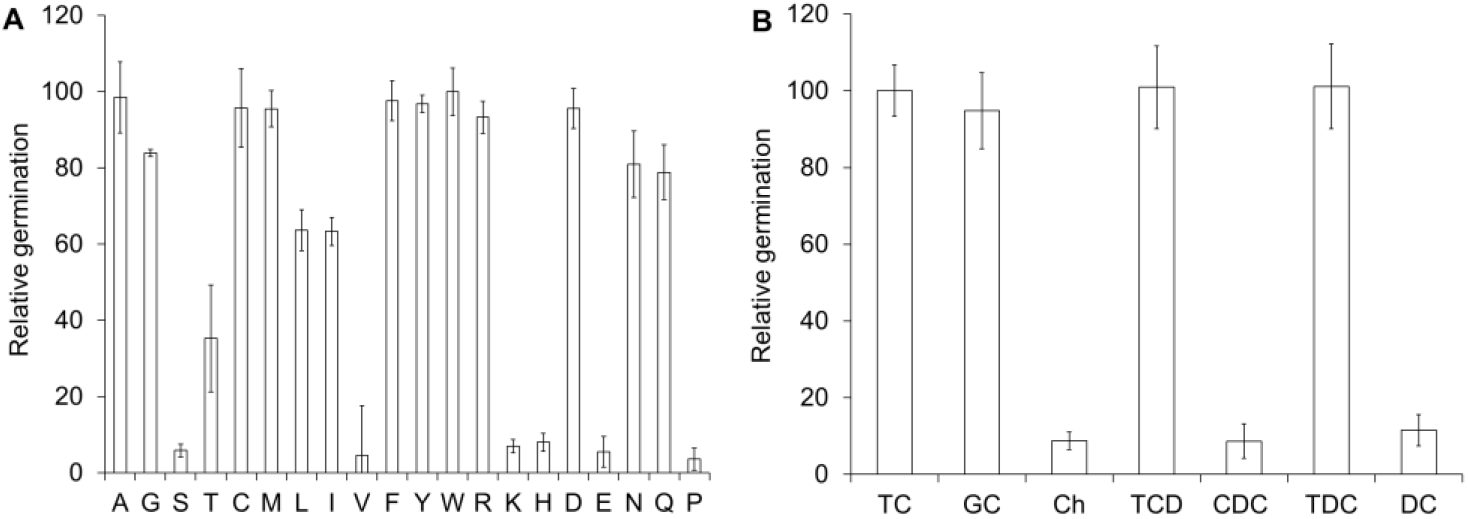
C. perfringens spores germinate with bile salts and amino acids. (A) *C. perfringens* JGS 1936 spores were treated with taurocholate and individual amino acids. Relative germination was calculated as fractions of germination rate for spores treated with L-alanine/taurocholate. Amino acids are represented by one-letter code. Error bars represents standard deviations of six independent measurements. (B) *C. perfringens* JGS 1936 spores were treated with L-alanine and either taurocholate (TC), glycocholate (GC), cholate (Ch), taurochenodeoxycholate (TCD), chenodeoxycholate (CDC), taurodeoxycholate (TCD), or deoxycholate (DC). Relative germination was calculated as fractions of germination rate for spores treated with L-alanine/taurocholate. Error bars represents standard deviations of six independent measurements.

Because bile salts serve to solubilize dietary fats [35], we treated *C. perfringens* spores with either SDS or Triton-X-100. Neither detergent was able to trigger *C. perfringens* spore germination even in the presence of excess L-alanine.

D-amino acids have been shown to inhibit amino acid-mediated spore germination in *Bacillus* species [36]. D-alanine and D-arginine failed to inhibit or induce *C. perfringens* spore germination in the AA germination pathway, but D-phenylalanine and D-tryptophan inhibited this pathway. In contrast, all the D-amino acids tested served as co-germinants with taurocholate in the BA pathway (data not shown).

### Optimization of *C. perfringens* spore germination conditions

To define the optimal conditions for *C. perfringens* spore germination, spores were germinated at different pH values. In the AA pathway, germination was significantly reduced if sodium phosphate buffer was substituted with potassium phosphate buffer (Fig. 3A). In contrast, the BA pathway was only active in the presence of potassium ions (Fig. 3B). For both pathways and in all strains, optimal germination occurred at near neutral to neutral pH. Germination was significantly reduced above pH 7.5 or below pH 5.5.

**Figure 3.**
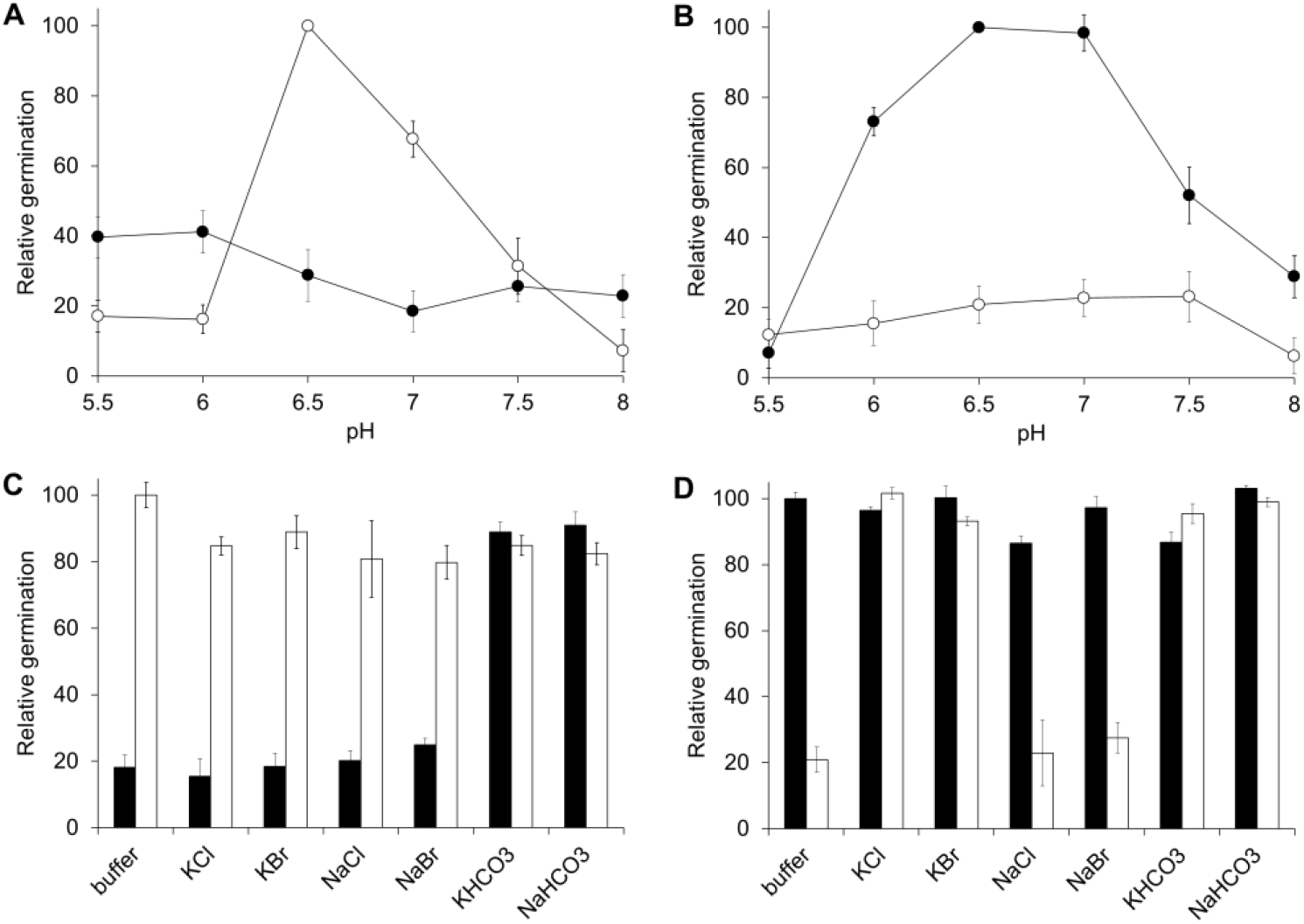
Effect of pH and ions on C. perfringens spores germination: (A) *C. perfringens* JGS 1936 spores were treated with L-alanine and L-phenylalanine at different pH values in sodium phosphate (○) or potassium phosphate (●) buffer. Relative germination was calculated as fractions of germination rate for spores suspended in sodium phosphate buffer, pH = 6.5. Error bars represents standard deviations of six independent measurements. (B) *C. perfringens* JGS 1936 spores were treated with taurocholate and L-alanine at different pH values in sodium phosphate (○) or potassium phosphate (●) buffer. Relative germination was calculated as fractions of germination rate for spores suspended in potassium phosphate buffer, pH = 6.5. (C) *C. perfringens* JGS 1936 spores were resuspended in either sodium phosphate (white bars) or potassium phosphate (black bars). Samples were then individually supplemented with KCl, KBr, NaCl, NaBr, KHCO_3_ or NaHCO_3_. Germination was initiated by addition of L-alanine and L-phenylalanine. Relative germination was calculated as fractions of germination rate for spores suspended in sodium phosphate buffer, pH = 6.5. (D) *C. perfringens* JGS 1936 spores were resuspended in either sodium phosphate (white bars) or potassium phosphate (black bars). Samples were then individually supplemented with KCl, KBr, NaCl, NaBr, KHCO_3_ or NaHCO_3_. Germination was initiated by addition of taurocholate and L-alanine. Relative germination was calculated as fractions of germination rate for spores suspended in potassium phosphate buffer, pH = 6.5.

Interestingly, addition of KCl, KBr NaCl, or NaBr did not affect the AA pathway response in either potassium phosphate or sodium phosphate buffer (Fig. 3C). Similarly, NaCl and NaBr did not affect the BA germination pathway when spores were resuspended in potassium phosphate or sodium phosphate buffer. On the other hand, addition of KCl or KBr induced the BA pathway in spores resuspended in sodium phosphate buffer (Fig. 3D).

Bicarbonate has been shown to be an essential co-germinant for some *Clostridium* species [18, 28]. In both the AA and BA germination pathways, addition of potassium bicarbonate or sodium bicarbonate increased germination rate for *C. perfringens* spores resuspended in both potassium and sodium phosphate buffers (Fig. 3C and 3D).

Because *C. perfringens* spores responded to germinants in a manner similar to *C. sordellii* and *C. difficile*, we tested all spore preparations by germination and growth in litmus milk medium. As expected for *C. perfringens*, all samples showed stormy clot fermentation [37]. The identities of selected spore samples were further confirmed by repeating 16S rRNA sequencing.

## Discussion and conclusions

*C. perfringens* is an animal pathogen with wide host distribution [25]. Pathogenic *C. perfringens* strains seem to be host specific [33]. To test if *C. perfringens* spores could impact host selection, we investigated the sporulation and germination profiles of pathogenic and non-pathogenic *C. perfringens* strains obtained from different hosts.

*C. perfringens* sporulation showed large strain variability. The source of peptone, the presence of theobromine, and the liquid media used for overnight bacterial growth contributed to this variability. Some strains sporulated under a single condition tested, while other strains had less stringent sporulation requirements. There was no apparent relationship between sporulation conditions and *C. perfringens* strain pathogenicity or host source. The best sporulation conditions for each strain are shown in Table S2.

Although *C. perfringens* sporulation conditions varied greatly between strains, the germination response was homogenous. Spores from different *C. perfringens* strains prepared under similar conditions behaved identically in germination assays. Furthermore, spores from a single strain prepared under different conditions were undistinguishable in germination assays. These results contrast with previous work suggesting that sporulation conditions affected the germination response of *Bacillus* spores [20-22].

Contrary to previous work, KCl, inosine, asparagine, glucose, fructose, sodium, and phosphate were unable to trigger *C. perfringens* spore germination [14, 17, 18]. The differences with these studies are not clear. We have previously shown that *Bacillus* spores purified by water washing alone show different germination profiles than spore preparations purified by gradient centrifugation [38]. Similar results were observed for *C. perfringens* spores. Water-washed spores were more heterogeneous and contained larger amounts of cellular debris. These contaminants can synergize with added metabolites to produce spurious germination responses not seen in cleaner spore preparation. Indeed, *C. perfringens* spore preparations purified by water washing auto-germinate upon storage. This heterogeneity will be more pronounce in *C. perfringens* spores due to the sensitivity of the germination response to the nature and order of addition of cations and bicarbonate.

Detailed analysis of germination conditions revealed two independent germination pathways (Fig. 4). The AA pathway required, at a minimum, L-alanine and L-phenylalanine to induce strong *C. perfringens* spore germination. L-arginine was not required but potentiates *C. perfringens* spore germination through the AA pathway. The competitive inhibition of the AA pathway by L-tryptophan, D-tryptophan, and D-phenylalanine suggests that binding of L-phenylalanine is a key process in germination triggering in this germination response.

**Figure 4.**
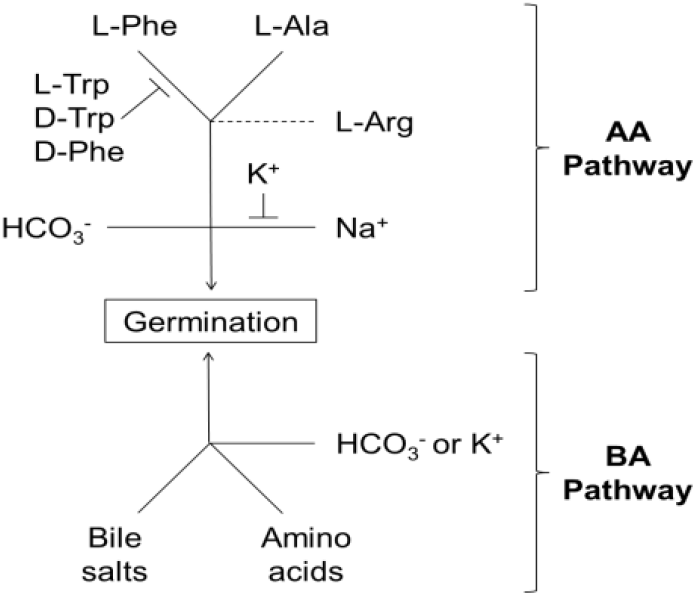
Scheme for C. perfringens spore germination: Solid lines represent required co-germinants. Capped lines represent germination inhibitors. Dashed lines represent germination enhancers.

The AA pathway also requires either bicarbonate or sodium ions. Sodium ions were necessary to activate the AA pathway in the absence of bicarbonate. Potassium acts as an inhibitor of the AA pathway, but only when added before sodium. We postulate that binding of cations is very fast and irreversible since once potassium is bound, sodium cannot induce germination of *C. perfringens* spores. Concomitantly, once sodium is bound, potassium is unable to inhibit germination. The presence of bicarbonate made the AA pathway insensitive to both potassium and sodium. This suggests that bicarbonate activates a separate, potassium insensitive, AA germination pathway.

In contrast to the limited number of amino acids used in the AA germination pathway, the BA germination pathway could be triggered by multiple combinations of amino acids and conjugated bile salts. The presence of salts also affected the BA germination pathway. Salt effect, however, was qualitatively different than in the AA pathway. Indeed, while L-tryptophan, D-tryptophan, and D-phenylalanine inhibited the AA pathway, these amino acids were strong co-germinants of the BA pathway. Potassium, an inhibitor of the AA pathway, acted as activator of the BA pathway. In contrast, sodium ions did not inhibit the BA pathway. Bicarbonate was also a strong activator of the BA pathway, bypassing potassium requirement. Neither pathway is activated or inhibited by phosphate, bromide or chloride.

*C. perfringens* spores share germination characteristics with both *C. sordellii* and *C. difficile*. The AA pathway was similar (but not identical) to the germination requirements of *C. sordellii* spores [28]. On the other hand, the BA germination pathway was similar (but not identical) the germination requirements of *C. difficile* spores [39, 40].

Recognition of bile salts by *C. perfringens* suggests that the sterane backbone is an ancestral germination signal as both *C. difficile* and *C. sordellii* recognize bile salts and progesterone analogs [34]. Whereas *C. sordellii* only uses steranes as enhancers of amino-acid mediated spore germination, both *C. difficile* and *C. perfringens* use bile salts as effective co-germinants. However, *C. perfringens* recognize a broader set of sterane germinants compared to *C. difficile*. More importantly, *C. perfringens* spores, similar to *C. sordellii*, are able to germinate using only amino acids as germination signals.

*C. perfringens* may have evolved multiple germination pathways to ensure spore response to different host environments. During the start of myonecrosis, *C. perfringens* spores would be exposed to abundant amino acid concentrations from decaying soft tissues. Furthermore, sodium and bicarbonate will be highly abundant in the fluids surrounding cells [41]. These are optimal conditions to activate the AA germination pathway (Fig. 4, top).

In enteric infections, however, *C. perfringens* spores would be exposed to the gastrointestinal tract environment. In the GI tract, bile salts and amino acids are abundant.

Furthermore, potassium concentrations are up to 20-fold higher than in serum [42]. These signals would favor using the BA germination pathway (Fig. 4, bottom) to establish infection.

## Acknowledgments

This work was supported by the National Institute of Food and Agriculture, U.S. Department of Agriculture, under Agreement No. 2010-65119-20603.

---

**Table S1.** Conditions for *C. perfringens* sporulation.

| **Table S1. Conditions for *C. perfringens* sporulation** |  |  |  |  |  |  |  |  |  |  |  |  |
| --- | --- | --- | --- | --- | --- | --- | --- | --- | --- | --- | --- | --- |
| Replated from BHI |  |  |  |  |  |  | Replated from Thioglycolate |  |  |  |  |  |
| Peptone number |  |  |  |  |  |  | Peptone number |  |  |  |  |  |
| Strain | #1 | #2 | #3 | #1* | #2* | #3* | #1 | #2 | #3 | #1* | #2* | #3* |
| 1936 | +++ | - | - | +++ | - | - | - | - | - | - | - | - |
| 1882 | - | ++ | - | - | +++ | - | - | - | - | - | - | - |
| 4064 | - | - | - | +++ | - | - | - | - | - | - | - | - |
| 1521 | - | - | - | - | - | - | - | - | - | +++ | - | - |
| 4151 | - | - | - | - | - | - | - | - | - | - | +++ | - |
| 1473 | - | - | +++ | - | - | + | ++ | - | - | +++ | - | - |
| 4104 | - | - | - | - | - | +++ | - | - | - | - | + | +++ |

All *C. perfringens* strains were plated on 2% agar supplemented with 1% yeast extract, 0.1% sodium thioglycollate, 60 mM Na<sub>2</sub>HPO<sub>4</sub>, 0.5% raffinose and 1.5% of a peptone source. \*Plates were also supplemented with theobromine to 0.01% final concentration.
+++ >40% spores. ++ 20-40% spores. + <20% spores. - no detectable spores

**Table S2.**
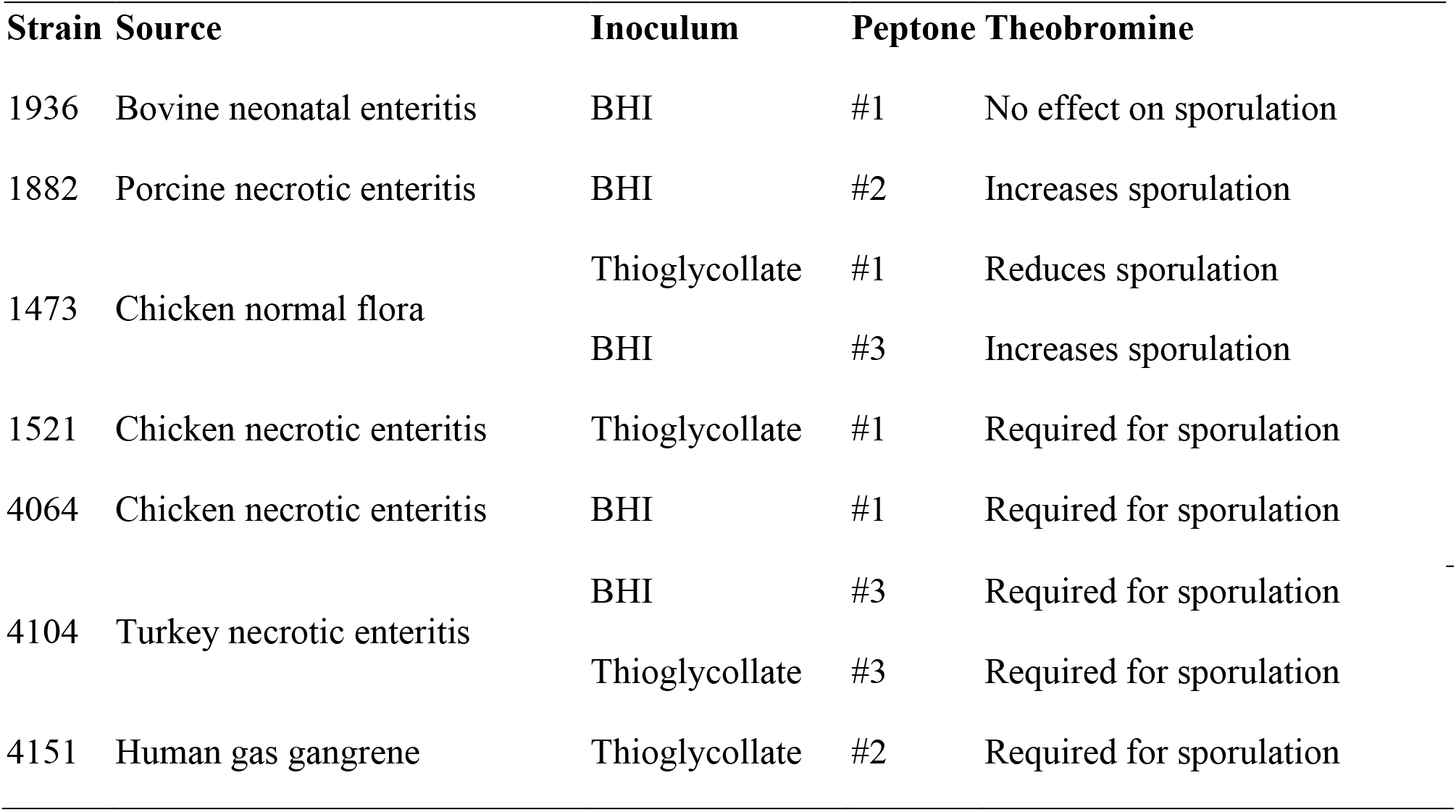
Source and optimal sporulation conditions for each *C. perfringens* strain.

